# Control over sampling boosts numerical evidence processing in human decisions from experience

**DOI:** 10.1101/2021.06.03.446960

**Authors:** Stefan Appelhoff, Ralph Hertwig, Bernhard Spitzer

**Affiliations:** Center for Adaptive Rationality, Max Planck Institute for Human Development

**Keywords:** active sampling, decision making, electroencephalography, information search, number processing

## Abstract

When acquiring information about choice alternatives, decision makers may have varying levels of control over which and how much information they sample before making a choice. How does control over sampling affect the quality of experience-based decisions? Here, combining variants of a numerical sampling task with neural recordings, we show that control over when to stop sampling can enhance (i) behavioral choice accuracy, (ii) the build-up of parietal decision signals, and (iii) the encoding of numerical sample information in multivariate electroencephalogram (EEG) patterns. None of these effects were observed when participants could only control which alternatives to sample, but not when to stop sampling. Furthermore, levels of control had no effect on early sensory signals or on the extent to which sample information leaked from memory. The results indicate that freedom to stop sampling can amplify decisional evidence processing from the outset of information acquisition and lead to more accurate choices.

## Introduction

Humans routinely acquire information about choice alternatives before deciding between them. In many situations, decision makers can control which and how much information they sample. For example, when deciding which of two products to buy, a customer may deliberately study reviews and testimonials before making a final choice. In other situations, the availability and amount of relevant information is determined by external factors. For instance, when selecting job applicants in an organization that uses standardized interviews, an employer must decide based on the applicants’ answers to the same set of predefined questions. More generally, decision scenarios can differ in the extent to which an agent has control over sampling, in terms of which and how much information is sampled before a choice is made.

One experimental setup suitable for studying how control over sampling may affect decision making is a numerical sampling paradigm (Hertwig et al. 2004; Hertwig and Erev 2009) in which participants can view sequential samples of possible choice outcomes before deciding for one or the other option. The paradigm has been used extensively in behavioral studies of risky choice to examine how decision makers choose between options they learned about from experience (i.e., through sampling the payoff distribution) as opposed to from formal description (e.g., “25% chance to obtain €10, otherwise €0”; Hertwig 2015; Wulff et al. 2018). Across these studies, researchers have also varied the extent to which participants were able to control the sampling process themselves. While the standard paradigm allows participants to decide freely which alternatives to sample and how often (Hertwig and Erev 2009), some studies have pre-specified the total number of samples to be taken (Hau et al. 2008; Ungemach et al. 2009; Fleischhut et al. 2014; Gonzalez and Mehlhorn 2016) or included matched (“yoked”) conditions in which participants had no control at all over the sampling sequence (Rakow et al. 2008). However, the latter variants of the sampling paradigm have been devised primarily to reduce confounds in comparisons with decisions from description (Rakow and Newell 2010); it remains unclear how control over sampling may alter experience-based decision making itself.

Several lines of evidence suggest that a sense of control can be beneficial in cognitive tasks (Gureckis and Markant 2012; Murayama et al. 2016). Agency in information acquisition has, for instance, been found to improve subsequent memory performance (Voss et al. 2011), even when exposure to the information was held constant (Murty et al. 2015). Another line of work has shown better performance in tasks self-selected by the participant than when the same tasks were selected by an experimenter (Murayama et al. 2015). More generally, various studies have identified performance benefits associated with volitional control per se and indicated that such effects could be mediated by motivational factors (Patall et al. 2008; Patall 2012). However, the effects of control cannot easily be generalized across domains. In several contexts, control does not seem to impact task performance (Flowerday and Schraw 2003; Flowerday et al. 2004) or can be detrimental—for instance, when control is perceived as irrelevant or as too complex (Katz and Assor 2007; but see Murayama et al. 2015). In the domain of decisions from experience using the sampling paradigm, evidence regarding the role of agency in the sampling process is still sparse. One recent meta-analysis suggested that control over sampling may alter the temporal weighting of numerical samples in subsequent choice (Wulff et al. 2018). However, this analysis was limited to comparisons across studies and did not address the performance benefits (or drawbacks) that may be associated with control over sampling, or the neurocognitive processes that might underlie them.

Here, we used specially designed variants of a numerical sampling paradigm combined with EEG recordings to study how control over sampling affects experience-based decision making. We systematically varied whether participants were free to decide which and how much information to sample (full control), or only which information to sample (partial control; with a pre-specified total number of samples), or whether they had no control over sampling at all. Critically, our design controlled for differences in stimulus presentation by matching the sample sequences in the no-control conditions with those in the self-controlled tasks. We found that full, but not partial control over sampling had a distinct beneficial effect on choice accuracy, and that this benefit was associated with a stronger encoding of numerical sample information from the outset of information acquisition.

## Materials and Methods

### Participants

Forty healthy volunteers took part in the experiment (20 female, 20 male; mean age 26.3 ± 3.7 years; all right-handed). All participants provided written informed consent and received a flat fee of €10 and €10 per hour as compensation, as well as a performance-dependent bonus (€9.35 ± €0.48 on average). The study was approved by the ethics committee of the Max Planck Institute for Human Development.

### Experimental design

On each trial, in all experimental conditions, participants were asked to decide between one of two choice options (left/right). Each choice option yielded one of two numerical outcomes (drawn from the range 1, 2, …, 9) with probability *p* (0.1, 0.2, …, 0.9), and the other outcome with probability 1 − *p*. The outcome values and probabilities on each trial were constrained such that (i) no two of the four numerical outcome values were identical and (ii) the difference in expected value between the two options was always 0.9 (derived from piloting). Under these constraints, the choice problems presented on each trial were selected pseudorandomly, with the additional restriction that each sample value (1, 2, …, 9) occurred with approximately equal probability across the experiment.

Half of the participants were assigned to the “full control” condition, where they were free to sample from the left or right option as often as they wished before making a final choice. The only restriction on sampling in the full-control condition was that a sample had to be taken within 3 seconds (otherwise the trial was restarted) and that the total number of samples could not exceed 19. The other half of participants were assigned to the “partial control” condition, which was identical to the full control condition except that a fixed number of 12 samples had to be drawn on every trial. The number of samples was based on pilot data where free-sampling participants took approximately 12 samples on average. In other words, participants in the partial control condition were also free to sample from the left or right option, but had no control over when to stop (or continue) sampling: They were always prompted to make a final choice after the 12th sample.

In both sampling conditions, the beginning of a new trial was signalled by a green fixation stimulus (a combination of bulls eye and cross hair; Thaler et al. 2013) that turned white after one second. Upon pressing the left or right button on a USB response pad (using the left or right hand respectively), participants were shown a black circular disk (diameter 5° visual angle) 4.5° to the left (choice option 1) or right (choice option 2) of fixation after 0.2 to 0.4 s (randomly varied). After another delay of 0.8 s, the number sample was presented in white (font Liberation Sans, height 4°) in the disk area for 0.5 s (see Figure 1 for a schematic illustration). After this, the disk disappeared and participants were given 3 s to draw the next sample. The black disk served as a spatial cue to minimize differences in surprise about the sample location (left/right) in yoked conditions without sampling control (see below). The sampling procedure was repeated depending on condition (partial control: 12 samples; full control: up to 19 samples), and the resulting sample sequences (including their precise timing) were recorded (see yoked conditions below). In the full control condition, a third button on the response pad (above the “right” button) was available to stop the sampling sequence. In all conditions, after the sampling was finished, the fixation stimulus changed color to blue for 1 s and participants were asked to make a final choice between the left and right options. The button and display procedure for the final choice was identical to that for drawing samples, except that the final choice outcome was displayed in green to indicate the eventually obtained reward. The rewards (i.e., the payouts from the final choices) were converted to Euros with a factor of 0.005 and added as a bonus to participant’s reimbursement after the experiment (see *Participants* above).

**Fig. 1.**
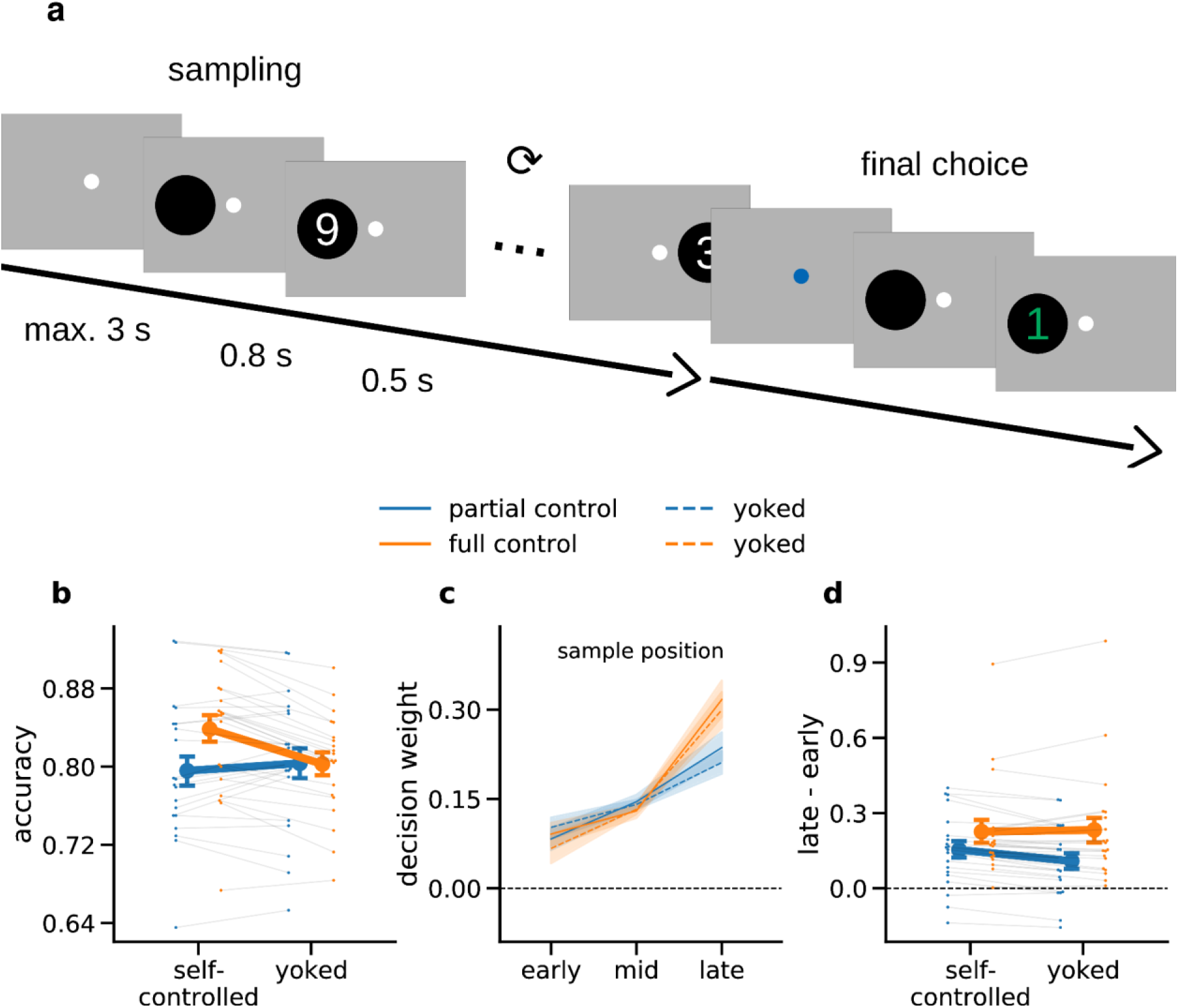
Experimental task and behavioral results. **(a)** Schematic illustration of an example trial. Participants were asked to decide between two choice options (left/right). Before committing to a choice, participants could draw up to 19 samples (full control group) or were required to draw a fixed number of 12 samples (partial control group). In yoked baseline conditions, participants judged replays of previously recorded sampling streams. **(b)** Mean accuracy (proportion of times the sampled sequence was judged correctly) in each condition. **(c)** Decision weights (see *Materials and Methods*) of samples occurring early, mid, or late in the sampling sequence, for each sampling condition. **(d)** Difference in decision weight between late and early samples. Higher values indicate that late samples had a stronger relative influence on choice than early samples (“recency” effect). Error indicators in all panels show SE.

Within both groups (full and partial control), each participant additionally performed the task in a “yoked” condition, where they had no control over sampling. Here, participants made decisions based on replays of previously recorded sampling streams (without any control over which and how many samples were shown or their timing). Accordingly, we refer to the yoked conditions as the no-control baseline conditions. In each group, half of the participants first performed the self-controlled sampling task (full or partial) and subsequently performed the no-control task with a replay of their own sampling sequences. In informal debriefing after the experiment, none of these participants reported having noticed that they had viewed exact replays of their own sampling sequences. The other half of the participants in each group performed the no-control task first (yoked to the sampling sequences of another participant in the same group) and the self-controlled task second. Control analysis showed no differences in choice accuracy between participants who performed the baseline task first (yoked to another participant’s sequences) or second (yoked to their own sequences) (all p > 0.05). Furthermore, in the subset of participants who were yoked to another participant, the difficulty of active versus yoked sampling sequences did not differ (all p > 0.05). Each participant performed 100 trials (five blocks of 20 trials with short breaks between blocks) in the self-controlled and yoked task variant, respectively.

Participants in the full control group drew on average 8.6 samples (SD = 4.2, median = 8), compared with the 12 samples that had to be drawn in the partial control group. Due to the principled impossibility of matching full and partial control trials (e.g., with respect to the precise length and timing of the sampling sequences on individual trials), all our analyses focus on comparisons of differences to the matched (yoked) baseline condition within each group. This analysis strategy rules out stimulus confounds that may arise, for instance, due to “amplification effects” under full control, where stopping decisions may be more likely when the momentary difference between the accumulated option values happens to be large (Hertwig and Pleskac 2010).

The experiment was programmed in Python using the Psychopy package (Peirce et al. 2019) and run on a Windows 10 PC. The experiment code is available on Zenodo (https://doi.org/10.5281/zenodo.3354368). Behavioral responses were recorded using a USB response pad (The Black Box ToolKit Ltd, UK). Throughout the experiment, eye movements were recorded using a Tobii 4C eye-tracker (Tobii Technology, Sweden; sampling rate 90 Hz). To reduce eye movements, participants’ gaze position was analyzed online while the experiment was run in all sampling conditions. The program displayed a warning message and restarted the trial whenever the gaze left an elliptical area centered on the central fixation stimulus (width 5° visual angle, height 2.85° visual angle) more than four times during a trial. Saccades towards the outcome samples were robustly detected with these settings. On average 3% of trials per participant were restarted due to lack of fixation, or failure to draw a sample within 3 seconds (see above). Offline analyses confirmed that participants generally held fixation in the remaining trials.

### Supplementary tasks

After the main experiment, participants performed an additional short task on the same choice problems, where the options were not explored through sampling but described formally on screen (e.g., “8 with 60% or 4 with 40%?”). Due to a coding error, much of the data (84%) from this task was incorrectly recorded and the results are thus not reported here. Participants further completed a brief numeracy questionnaire (Berlin Numeracy Test, BNT (Cokely et al. 2012)). Exploratory analysis showed no significant correlations of the effects reported in our main analysis with BNT scores.

### EEG recording

The experiment was performed in an electrically shielded and soundproof cabin. Scalp EEG was recorded with 64 active electrodes (actiCap, Brain Products GmbH Munich, Germany) positioned according to the international 10% system. Electrode FCz was used as the recording reference. We additionally recorded the horizontal and vertical electrooculogram (EOG) and electrocardiogram (ECG) using passive electrode pairs with bipolar referencing. All electrodes were prepared to have an impedance of less than 10kΩ. The data were recorded using a BrainAmp DC amplifier (Brain Products GmbH Munich, Germany) at a sampling rate of 1000 Hz, with an RC high-pass filter with a half-amplitude cutoff at 0.016Hz (roll-off: 6dB/octave) and low-pass filtered with an anti-aliasing filter of half-amplitude cutoff 450Hz (roll-off: 24dB/octave). The dataset is organized in Brain Imaging Data Structure format (BIDS; Gorgolewski et al. 2016) according to the EEG extension (Pernet et al. 2019), and is available from https://gin.g-node.org/sappelhoff/mpib_sp_eeg/.

### Behavioral data analysis

Participants’ behavioral accuracy in each sampling condition was calculated with respect to the arithmetic mean of the samples that were presented for each option. Differences in accuracy between sampling conditions were analyzed using a mixed 2×2 ANOVA (self-controlled/yoked; full/partial), followed up with Bonferroni-corrected pairwise t-tests. All statistical tests reported (including in the EEG analyses, see below) are two-tailed.

To examine recency effects in the behavioral data, we used a reverse correlation approach (Neri et al. 1999; Spitzer et al. 2016) based on logistic regression. We first divided the samples in a trial into early, mid, and late samples. The first and last two samples in a trial were defined as early and late samples, respectively, and the remaining samples as “mid” samples. Trials with fewer than 5 samples overall were discarded in this analysis (between 1% and 41.5% of trials per participant, mean = 13.3%). For each participant, task condition, and time window, we regressed the participant’s final choices (left: 0, right: 1) onto the number sample values (numbers 1, 2, …, 9 rescaled to −4, −3, …, 4), where the values for the left option were sign-flipped to reflect their opposite impact on the probability of choosing the right option (Spitzer et al. 2017). We interpret the regression coefficients resulting from this analysis as a measure of “decision weight”, that is, of the influence that number samples (early, mid, or late) had on choice.

### EEG preprocessing

The EEG recordings were visually inspected for noisy segments and bad channels. Ocular and cardiac artifacts were corrected using independent component analysis (ICA). To this end, we high-pass filtered a copy of the raw data at 1 Hz and downsampled it to 250 Hz. We then ran an extended infomax ICA on all EEG channels and time points that were not marked as bad in the prior inspection. Using the EOG and ECG recordings, we identified stereotypical eye blink, eye movement, and heartbeat artifact components through correlation with the independent component time courses. We visually inspected and rejected the artifact components before applying the ICA solution to the original raw data (Winkler et al. 2015). We then filtered the ICA-cleaned data between 0.1 and 40 Hz, interpolated bad channels, and re-referenced each channel to the average of all channels. Next, the data were epoched from −0.2 to 0.8 s relative to each individual number sample onset. Remaining bad epochs were rejected using a thresholding approach from the FASTER pipeline (Step 2; Nolan et al. 2010). On average, n = 1925 clean epochs (93.85%) per participant were retained for analysis. The epochs were downsampled to 250 Hz and baseline corrected relative to the period from −0.2 to 0 s before stimulus onset. All EEG analyses were performed in Python using MNE-Python (Gramfort et al. 2013), MNE-BIDS (Appelhoff et al. 2019), and custom code. All analysis code is available at https://github.com/sappelhoff/sp_code.

### Event-related potential (ERP) analysis

EEG analyses are reported for the epochs around the onset of the individual number samples. We first examined lateralized visual ERP components to test whether early visual processing differed between the sampling conditions. To this end, we subtracted the ERP for stimuli presented on the right from the ERP for stimuli presented on the left, and then subtracted the mean signal of right-hemispheric (O2, PO4, PO8, PO10) occipito-parietal channels of interest (based on previous literature; Eimer 1998) from the corresponding left-hemispheric (O1, PO3, PO7, PO9) channels. Mean amplitudes of the lateralized evoked potential were extracted from prototypical time windows (P1 ERP component: 80 ms to 130 ms, N1 ERP component: 140 ms to 200 ms) for each sampling condition and analyzed in a mixed 2×2 ANOVA (self-controlled/yoked; full/partial).

We further examined centro-parietal evoked responses (CPP/P3, averaged over the early, mid, and late samples in each trial) as a potential correlate of decisional evidence accumulation (O’Connell et al. 2012; Twomey et al. 2015; Pisauro et al. 2017). To this end, we averaged the signal over centro-parietal channels (Cz, C1, C2, CPz, CP1, CP2, CP3, CP4, Pz, P1, P2) and focused on a time window from 300 ms to 600 ms, based on previous analyses of CPP/P3 responses during visual stimulus sequences (Spitzer et al. 2017; Wyart et al. 2015; Polich 2007).

### Representational similarity analysis (RSA)

To examine the encoding of numerical sample value in multivariate EEG patterns, we used an approach based on representational similarity analysis (RSA; Kriegeskorte and Kievit 2013). To this end, the ERPs were additionally smoothed (Grootswagers et al. 2016) with a Gaussian kernel (35 ms half duration at half maximum). We then computed multivariate (dis)similarity in terms of the pairwise Euclidean distance between the ERP patterns associated with each sample value (numbers 1–9), yielding a 9×9 representational dissimilarity matrix (RDM) for each time point of the analysis epoch. The EEG RDMs at each time point were compared with model RDMs reflecting (1) the samples’ numerical magnitude (“numerical distance”; upper panel Fig. 3a) and (2) their “extremity” (i.e., a sample’s absolute difference from the midpoint of the sample range, 5; Fig. 3d). To avoid confounds by potential deviations from a uniform distribution of sample values across the experiment, we additionally orthogonalized each model RDM to an RDM of the relative frequency of numerical sample occurrences (Spitzer et al. 2017). Qualitatively similar results were obtained when this orthogonalization step was omitted.

For quantitative analysis, we extracted the lower triangle (excluding the diagonal) from each model RDM and compared it with the EEG RDM data at each time point using the Pearson correlation coefficient. To analyze the overall encoding of numerical distance and extremity, we used t-tests against zero with cluster-based permutation testing (Maris and Oostenveld 2007) to control for multiple comparisons over time points (10,000 iterations, cluster-defining threshold = 0.05). We then re-computed the EEG RSA separately for each sampling condition to test for differences in number encoding. Differences between conditions were examined using mixed 2×2 ANOVAs (self-controlled/yoked; full/partial), again using cluster-based permutation testing to control for multiple comparisons over time points. Analogous RSA analyses were performed separately on the first and second half of samples from each trial (Fig. 3c).

## Results

Participants (n = 40) observed sequential samples (Arabic digits 1–9) of the potential payouts of choice options (left/right) before deciding on one of them (Fig. 1a). In different conditions, participants (i) could determine from which option(s) to sample and when to stop sampling (“full control”, 1–19 samples/trial, n = 20 participants) or (ii) could determine only from which option to sample for a fixed number of samples (“partial control”, 12 samples/trial, n = 20 participants). Each participant additionally performed the task in a “yoked” condition with matched sample sequences (see *Materials and Methods*) that they could not control. Our behavioral and EEG analyses focus on the effects of control (full or partial) relative to the matched (yoked) no-control conditions.

### Behavior

We examined choice accuracy with respect to the average value of the samples observed in each option. Mean choice accuracy was 83.8% under full control (SE = 1.4%, yoked baseline: 80.3%, SE = 1.2%) and 79.6% under partial control (SE = 1.6%, yoked baseline: 80.3%, SE = 1.6%). A mixed 2×2 ANOVA with the factors control over sampling (self-controlled or yoked; within participants) and control type (full or partial; between participants) showed no main effects [self-controlled/yoked: F(1,38) = 2.143, p = 0.151, η_p_^2^ = 0.053; full/partial: F(1,38) = 1.321, p = 0.258, η_p_^2^ = 0.034], but a significant interaction of the two factors [F(1,38) = 5.108, p = 0.03, η_p_^2^ = 0.118)]. Post hoc tests showed significantly higher accuracy under full control than in the yoked baseline [t(19) = 2.644, p = 0.032, d = 0.605, Bonferroni corrected], but no such effect under partial control [t(19) = −0.561, p > 0.9, d = −0.108]. Thus, relative to matched baseline conditions, we found an accuracy benefit of control over sampling under full control, but not under partial control.

We next examined if and to what degree the integration of sample information over time differed between conditions. To this end, we examined the samples’ decision weights (see *Materials and Methods*, *Behavioral data analysis*) separately for early, mid-, and late portions of the sampling sequence (Fig. 1c). We found a pronounced “recency” pattern (Tsetsos et al. 2012; Cheadle et al. 2014), with decision weight increasing over the course of the trial. In other words, later samples generally had a higher impact on the final choice than earlier samples. For comparison between sampling conditions, we quantified recency as the difference in decision weight between late and early samples (Fig 1d). A mixed 2×2 ANOVA, specified analogously as for accuracy above, showed no significant main effects [self-controlled/yoked: F(1,38) = 0.8, p=0.377, η_p_^2^ = 0.021; full/partial: F(1,38) = 3.363, p = 0.075, η_p_^2^ = 0.081], and no interaction between the two factors [F(1,38) = 1.483, p = 0.231, η_p_^2^ = 0.038]. Thus, we found no impact of control over sampling on recency. Together, full control over sampling was characterized by increased choice accuracy but was not distinguished in the extent to which sample information “leaked”(Usher and McClelland 2001), or was forgotten, in the course of a trial.

### Visual evoked responses

Turning to the EEG data, we first examined visual evoked responses to test whether the sampling conditions differed in terms of early sensory processing of the sample stimuli (e.g., due to potential differences in stimulus-directed visual attention; Luck et al. 2000). Figure 2a shows the occipitoparietal ERP difference between stimuli occurring in the right and left visual fields, subtracted between contralateral channels (see *Materials and Methods*, *ERP analysis*). Statistical analysis showed no differences between sampling conditions in the time window of either the P1 (80–130ms) or the N1 component (140–200ms) of the visual ERP [all F(1,38) < 1.71, all p > 0.20, all η_p_^2^ < 0.044; mixed 2×2 ANOVAs specified as in the behavioral analysis above]. We thus found no evidence for differences in early visual processing between the sampling conditions.

**Fig. 2.**
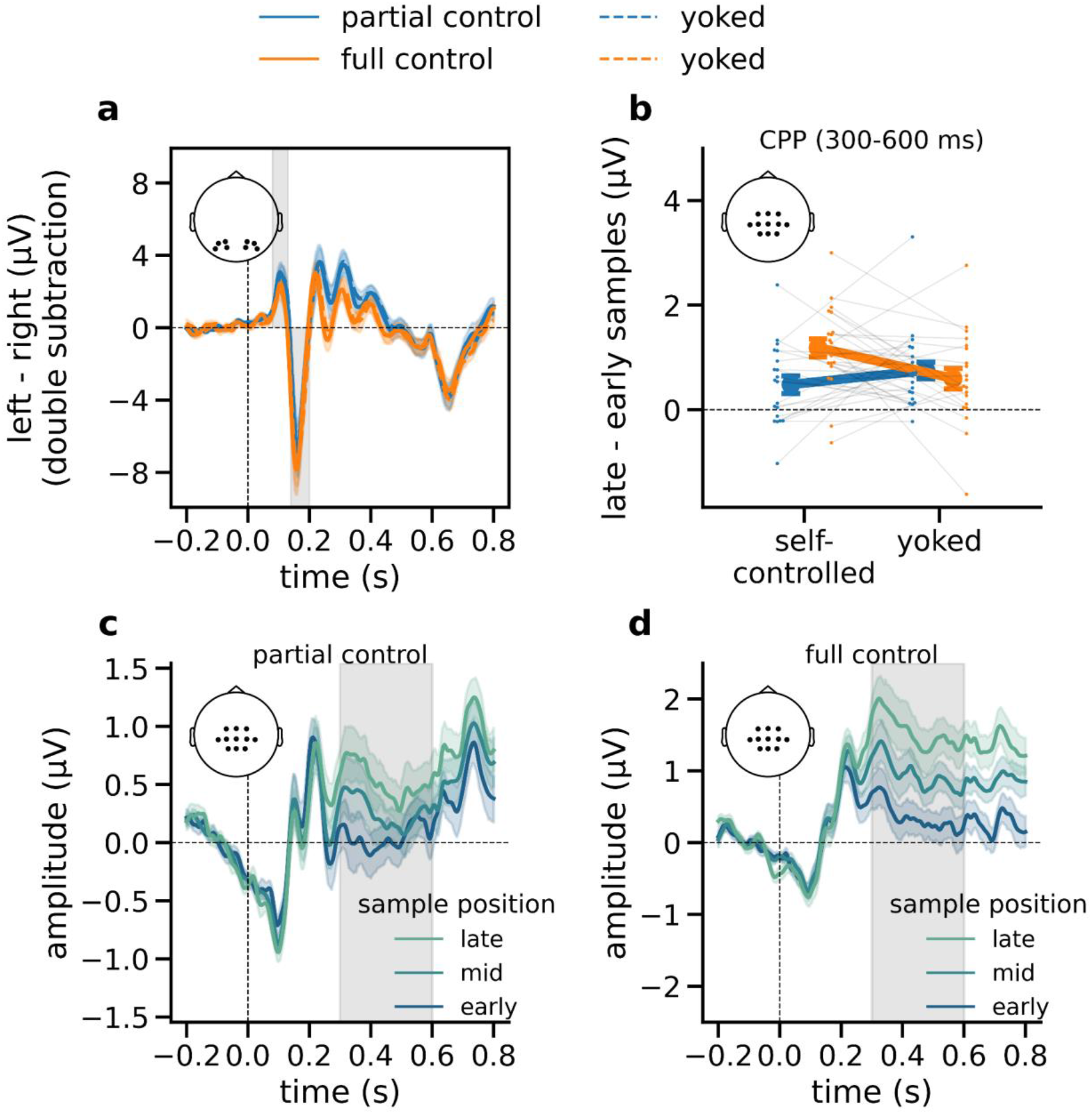
Univariate EEG results with ERPs time-locked to number sample onset. **(*a)*** Early visual ERPs (left − right stimuli, right channels subtracted from left channels) in each sampling condition. Gray shadings indicate time windows of the P1 and N1 components, respectively (80–130 ms and 140–200 ms). **(b)** The difference in centro-parietal (CPP) amplitudes between samples occurring late vs. early in the trial (see panels c–d), plotted separately for each sampling condition (including yoked). **(c)** The “ramping up” of CPP amplitudes over early, mid, and late samples in the partial control condition. Gray shadings indicate the time window from which average amplitudes were extracted in panel *b*. **(d)** same as *c*, for the full-control condition. Error indicators in all panels show SE.

### Centro-parietal positivity (CPP) / P3

We next examined CPP responses over centro-parietal channels between 300 ms and 600 ms after stimulus onset. The amplitude of the CPP response to a sample generally increased in the course of the trial (Figures 2b–d), which is in line with previous studies implicating the CPP in decisional evidence accumulation (O’Connell et al. 2012; Twomey et al. 2015). Figures 2c and 2d illustrate the monotonic ramping-up of CPP across samples occurring early, mid, and late in the trial (see *Materials and Methods*) under partial and full control. Descriptively, the build-up of CPP was stronger under full control. For statistical analysis, we examined the increase in CPP amplitude from early to late samples in the individual sampling conditions (Fig. 2b). A significant increase in amplitude was evident in each condition (including yoked; Fig 2b, all p < 0.02, t-tests against zero, uncorrected). A mixed 2×2 ANOVA comparing the amplitude difference between conditions showed no significant main effects [self-controlled/yoked: F(1,38) = 1.579, p = 0.217 η_p_^2^ = 0.04; full/partial: F(1,38) = 1.534, p = 0.223, η_p_^2^ = 0.039], but a significant interaction [F(1,38) = 11.408, p = 0.002, η_p_^2^ = 0.231]. Post-hoc t-tests showed that the CPP increased more steeply in the full control condition than in the yoked baseline [t(19) = 2.772, p = 0.024, d = 0.687, paired t-test, corrected], whereas no such effect was evident under partial control [t(19) = −1.932, p = 0.137, d = −0.355]. These findings suggest that the increased choice accuracy under full control was associated with a stronger build-up of a cumulative decision variable (Twomey et al. 2015), as indicated by a steeper increase of centro-parietal decision signals within trials.

### RSA

Our results so far show that decisions made with full control over sampling were more accurate and accompanied by a stronger build-up of parietal choice signals (Fig. 2c–d), but there was no evidence for differences in early visual processing (Fig. 2a) or in the leakage of sample information over time (Fig. 1b–c). One possibility is that a benefit of full control may have arisen at the stage of numerical processing, in encoding the abstract decisional value of the sampled number information proper. We used an RSA-based approach (see *Materials and Methods*) to examine the neural encoding of numerical sample values, building on previous findings of numerical distance effects in multivariate EEG patterns (Spitzer et al. 2017; Teichmann et al. 2018; Luyckx et al. 2019; Sheahan et al. 2021). Specifically, we correlated the multivariate similarity structure of samples (1–9) in our EEG data with theoretical models reflecting (i) numerical distance and (ii) extremity of the sample values (see *Materials and Methods*).

#### Numerical distance

We found robust encoding of numerical distance in multivariate EEG signals between approximately 160 and 800 ms after sample onset (Fig. 3b, p_cluster_ < 0.001, t-test against zero), replicating previous findings in tasks without sampling control (Spitzer et al. 2017; Teichmann et al. 2018; Luyckx et al. 2019; Sheahan et al. 2021). To test whether the strength of this effect differed between levels of sampling control, we examined its time course in the various conditions (full, partial, yoked baselines) using mixed 2×2 ANOVAs (specified analogously as above). The analysis showed no main effects (all p_cluster_ > 0.05) but a significant interaction cluster between 320 ms and 580 ms (p_cluster_ = 0.009). We further compared the average numerical distance effects in the time window of this cluster. We found the effect to be significantly larger (relative to yoked baseline) under full control [t(19) = 3.65, p = 0.003, d = 1.05, corrected], but not under partial control [t(19) = −1.065, p = 0.6, d = −0.340, corrected]. In other words, the encoding of number information in sample-level neural signals was enhanced under full control, mirroring the pattern of findings for CPP build-up (Fig. 2b) and choice accuracy (Fig. 1b).

**Fig. 3.**
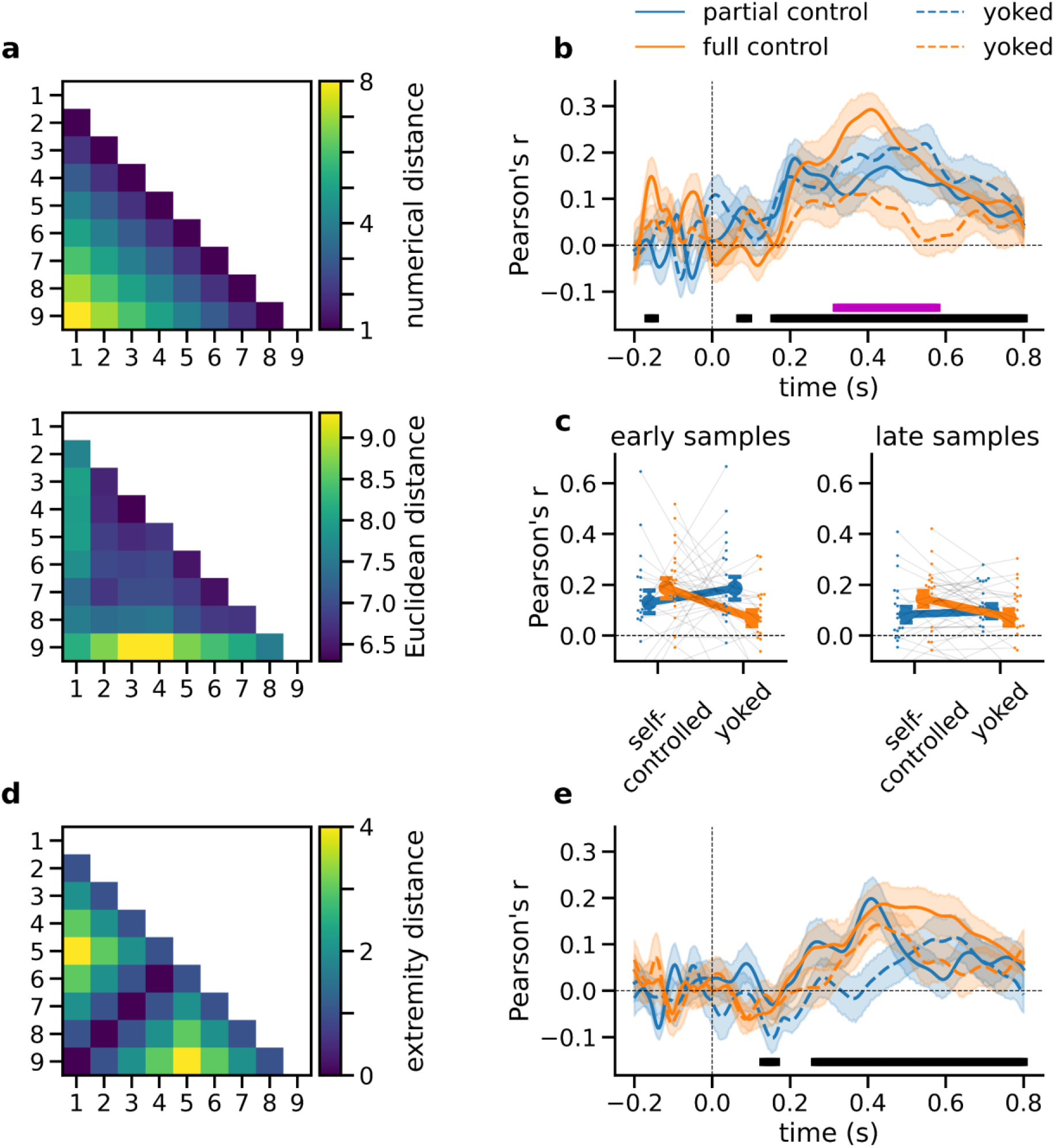
RSA Results. **(a)** Upper: Model RDM reflecting the pairwise numerical distance between sample values. Lower: Grand mean EEG RDM averaged across participants and sampling conditions in a representative time window between 300 and 600 ms after sample onset. **(b)** Time course of numerical distance effects in multivariate EEG patterns, plotted separately for each sampling condition. Black bar indicates time windows of significant numerical distance encoding (collapsed across sampling conditions). Purple bar indicates the time window of significant differences between sampling conditions (interaction effect, see *Results*). **(c)** Mean numerical distance effects by condition. Left: First half of samples in each choice trial. Right: Second half. **(d)** Model RDM reflecting the sample values’ extremity in terms of their absolute distance from the midpoint of the sample range (i.e., 5). **(e)** Time course of extremity encoding in multivariate EEG, plotted separately for each sampling condition. All error bars and shadings show SE.

We next asked whether the enhanced number encoding under full control was driven solely by late samples occurring near the time of the decision to stop sampling. To this end, we repeated the RSA analysis separately for the first (Fig. 3c, left) and second (Fig. 3c, right) half of the samples in a trial. Importantly, a significant enhancement under full control was already evident in the first half of samples [t(19) = 2.279, p = 0.034, d = 0.707], that is, long before participants stopped sampling. The effect in the second half of samples was similar [t(19) = 2.237, p = 0.037, d = 0.673; partial control: both p > 0.24]. In sum, we found no indication that enhanced number encoding under full control occurred only near the time of deliberate (vs. forced) stopping. Rather, the effect appeared to emerge early in the sampling sequence. We note again that we only interpreted effects in relation to the respective matched (yoked) control conditions, as other comparisons may suffer from non-trivial confounds (see *Materials and Methods*, *Experimental design*).

#### Extremity

Inspection of the empirically observed RDM (Fig. 3a, lower) suggests that besides numerical distance, the multivariate EEG patterns also encoded the extremity of the sample values (i.e., their absolute distance from the mid-point of the sample range (Spitzer et al. 2017; Luyckx et al. 2019)). Using a model RDM of numerical extremity (Fig. 3d; note that the model is orthogonal to the numerical distance RDM in Fig. 3a, upper), we found a significant effect between approximately 260 ms and 800 ms (t-test against zero, p_cluster_ < 0.001) in the EEG data collapsed across conditions. However, testing for differences between sampling conditions yielded no significant results (all p_cluster_ > 0.05). Together, while both numerical distance and numerical extremity were reflected in our multivariate EEG data, only numerical distance mirrored the enhancement under full control that was observed in CPP build-up and in behavior.

#### Neurometric distortions

Recent studies of sequential number comparisons (without participant control over sampling) have shown that neural number representations can be distorted (e.g., anti-compressed) away from the linearly monotonic distance structure of our idealized model RDMs (Fig. 3a). We used a “neurometric” approach (Spitzer et al. 2017) to test (i) whether such distortions were replicated in our task and (ii) whether they differed between levels of control. To this end, we parameterized our model RDMs to reflect the distance structure of transformed values *ν* = *sign*(*x* + *b*) |*x* + *b*|^*k*^, where *x* are the numerical sample values (1–9 normalized to the range [−1, 1]), exponent *k* determines the shape of the transformation (*k* < 1 compression; *k* = 1 linear; *k* > 1 anti-compression), and *b* reflects a bias towards smaller (*b* < 0) or larger numbers (*b* > 0). Our EEG data, averaged across all conditions, were best explained by parameterizations *k* > 1 and *b* > 0 (Fig. 4a; both p < 0.003, t-tests of individual subject maxima against 1 and 0, respectively, averaged over parameterized distance and extremity). Thus, the neural number representation was anti-compressed and biased towards larger magnitudes (Fig. 4b), strongly resembling the distortions observed in previous work (Spitzer et al. 2017; Luyckx et al. 2019). In comparisons between levels of control, however, we found no evidence for differences in the degree of anti-compression (Fig. 4c, left; both p > 0.545, t-tests of *k* against yoked baselines or bias; Fig. 4c, right; both p > 0.131, t-tests of *b* against yoked baselines). In other words, under full control, the encoding of numerical sample information was amplified (Fig. 3) without any notable changes in its general representational geometry.

**Fig. 4.**
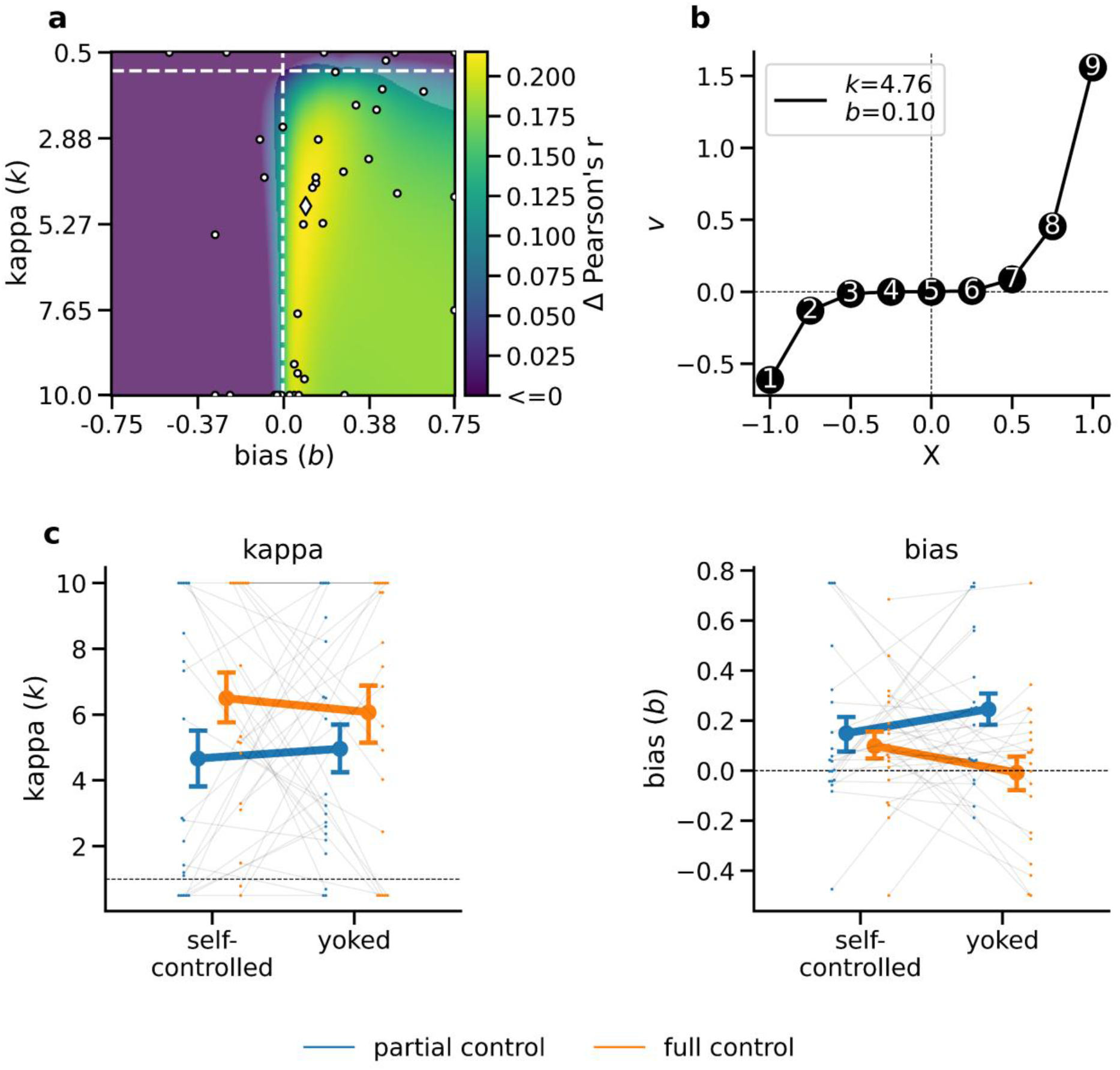
Neurometric distortions. **(a)** Grand mean neurometric map, combined across all task conditions. Color scale indicates change in EEG encoding strength (*Δ r*, averaged over distance- and extremity models) as a function of non-linear distortions of numerical value (*k* < 1: compression; *k* > 1: anti-compression, *b*: bias). Dashed lines indicate linear (*k* = 1) and unbiased (*b* = 0) models. Parts of the map that are not overlaid with a slightly opaque mask contain values with a significant increase relative to unbiased linear encoding (p < 0.001, corrected using false discovery rate). White markers show maxima (diamond: mean; dots, individual participants). **(b)** Neurometric function, parameterized according to the maximum mean correlation identified in *a*. **(c)** Neurometric parameter estimates in the individual sampling conditions, *left*: exponent (*k*); *right*: bias (*b*). Error bars show SE.

## Discussion

Using variants of a numerical sampling paradigm and controlling for stimulus confounds, we observed increased choice accuracy when participants had control over the sampling process before committing to a choice. On the neural level, the behavioral benefit was reflected in a stronger encoding of the numerical sample information in multivariate EEG patterns and in a steeper build-up of centroparietal choice signals. The key determinant of these effects was participants’ control over *how much* information to sample. Freedom to decide only which options to sample, but not when to stop sampling, did not bring about the same effects, in either behavior or neural signals.

Drawing on a well-established sequential sampling framework (Gold and Shadlen 2007; Ratcliff and McKoon 2008; O’Connell et al. 2012), our behavioral and neural findings provide a neurocognitive perspective on how control over sampling may boost choice accuracy. We observed no differences in early visual ERPs known to be modulated by top-down visual attention (Mangun and Hillyard 1991; Luck et al. 1994, 2000), but a robust enhancement further downstream in the processing hierarchy, at the level of symbolic number encoding (Ansari et al. 2005; Nieder and Dehaene 2009). Our results replicate recent findings of a “neuronal numberline” in multivariate EEG patterns, where the neural representation of for example, number “6” is more similar to that of “7”, than to that of “9” (Spitzer et al. 2017; Teichmann et al. 2018; Luyckx et al. 2019; Sheahan et al. 2021). We found this representation of numerical magnitude to be amplified under full control, mirroring the pattern observed in behavioral performance. Importantly, number encoding was already enhanced for samples occurring early in the trial, long before participants stopped sampling to make a final choice. Consistent with this finding, we also observed a steeper rise in parietal indices of evidence accumulation (CPP/P3; O’Connell et al. 2012; Twomey et al. 2015) *across* samples, as if each individual sample contributed stronger evidence to the ongoing decision formation. In a sequential sampling framework where evidence is accumulated into a running decision variable (Gold and Shadlen 2007; Kiani et al. 2008; Ratcliff and McKoon 2008; O’Connell et al. 2012; Glickman and Usher 2019), our EEG and behavioral findings may thus both be attributable to an improvement in numerical evidence processing.

One possible explanation for our findings relates to motivational factors. Previous work has shown that the ability to actively control the environment and/or one’s subjective experiences can have beneficial effects, for example on memory (Voss et al. 2011; Murty et al. 2015), self-regulation and error monitoring (Legault and Inzlicht 2013), learning and inductive inference (Gureckis and Markant 2012; Markant and Gureckis 2014), and various other aspects of cognition and behavior (Patall et al. 2008; Leotti et al. 2010; Leotti and Delgado 2011; Patall 2012; Murayama et al. 2016). Our findings add to these literatures by showing that control can also confer benefits in sample-based decision making, specifically when participants can control when to stop sampling. While the extrinsic rewards for choice accuracy were identical across our task conditions, control over stopping can add an incentive to optimize the time spent on a trial (Ostwald et al. 2015; Tickle et al. 2020). There is typically a trade-off between speed and accuracy of task execution (Heitz 2014), such that faster decisions come at the cost of lower accuracy (but see Gigerenzer et al. 2011). However, the present findings under full control cannot be explained by such a trade-off, given that we observed benefits relative to yoked trials of identical length. As we used exact copies of the participant-generated sampling sequences in our baseline conditions, we can also rule out the possibility that the results are attributable to “amplification effects” (Hertwig and Pleskac 2010), where participants tend to stop sampling when the cumulative difference between options happens to be large (leading to objectively easier trials; see below). With these simpler explanations ruled out, our findings suggest that control per se may lead to more efficient sample encoding, potentially through increased task engagement when decision time can be optimized on a trial by trial basis.

We found no differences between conditions in the temporal weighting of sample information over the course of a trial. A clear recency effect (relative overweighting of late samples) was evident in all task conditions, including yoked baselines. This pattern appears to be at odds with a previous meta-analysis of numerical sampling studies (Wulff et al. 2018), where recency effects were observed solely in conditions with full agency over sampling. However, the present pattern is consistent with established findings of recency effects in other sequential integration tasks where sample presentation is entirely experimenter-controlled (Tsetsos et al. 2012; Cheadle et al. 2014; Wyart et al. 2015; Spitzer et al. 2017; Glickman and Usher 2019; Luyckx et al. 2019; Kang and Spitzer 2021). We found no evidence that the extent to which sample information presented early in the sequence is down-weighted (or “leaks” from memory; Usher and McClelland 2001) was affected by control over sampling. We also found no indication of differences in the representational geometry of the sampled information in neural signals. Neurometric analysis showed an anti-compression of numerical values (Spitzer et al. 2017; Luyckx et al. 2019) in all conditions, regardless of the level of control. The absence of differences in these more qualitative aspects of information processing in our tasks suggests that the cognitive benefits of full control may best be described as an overall increase in the gain of neural processing (Donner and Nieuwenhuis 2013; Eldar et al. 2013; Murphy et al. 2016), which amplify the critical decisional information in a sample (here, numerical magnitude).

None of the benefits observed under full control were evident in our partial control condition, where participants could only decide which option to sample next, but not when to terminate sampling. Although the partial control condition gave participants some level of agency (relative to the yoked conditions without control; Chambon et al. 2020; Weiss et al. 2021), we suspect that it may not have induced a strong sense of control over the task. It even seems possible that participants may have perceived the requirement to perform a prescribed number of sampling actions as externally controlled and a cognitive burden (see also Sullivan-Toole et al. 2017). Indeed, post hoc examination of left/right sampling patterns showed that our participants resorted to stereotypical sampling routines (either alternating between options: “a-b-a-b-…” or sampling first one option and then the other: “a-a-a-…-b-b-b”) in 67.61% of trials (relative to the yoked conditions without control; for related findings see Hills and Hertwig 2010). In other words, participants made little use of the freedom to vary their left/right sampling strategy trial by trial (and/or sample by sample), potentially due to a lack of perceived benefits (Dixon and Christoff 2012). In this light, it is perhaps not surprising that we found no processing enhancements under partial control, in either behavior or neural signals.

Numerical sampling tasks similar to ours have been used extensively in the past to study decisions from experience (Hertwig et al. 2004) in complement to the common use of symbolic descriptions to study risky choice (Kahneman and Tversky 1979; Juechems et al. 2021). Experience-based choices can differ systematically from description-based choice, especially in terms of probability weighting (Hertwig and Erev 2009; Wulff et al. 2018). A much-discussed aspect of this “description–experience gap” is that participants in experience-based tasks tend to rely on relatively few samples (Hau et al. 2010; Plonsky et al. 2015; Wulff et al. 2018). Also in our experiment, participants in the full-control condition chose to sample less than they could have (Furl and Averbeck 2011). Although one explanation is that small samples can render choices objectively simpler (Hertwig and Pleskac 2008, 2010), our findings suggest that small samples may also defy typical accuracy trade-offs if the decision to stop sampling lies in the autonomy of the sampling agent (see also Petitet et al. 2021). Granting participants full control over sampling may thus not only enable but directly promote reliance on small samples through more efficient processing of the sample evidence.

Finally, next to an encoding of the numerical sample information in multivariate EEG patterns, we also found an apparent encoding of “extremity” (Fig. 3d–e), which did not differ between levels of control over sampling. Future research may further investigate this effect with regards to extreme events in decisions from experience (Ludvig and Spetch 2011; Ludvig et al. 2014, 2018).

In summary, we found that control over sampling can enhance the neural encoding of decision information and improve choice accuracy. The results add to a growing collection of findings that exercising agency can benefit performance in cognitive tasks and shed light on the neural processes that support such benefits.

## Data availability

All data in BIDS format are available at https://gin.g-node.org/sappelhoff/mpib_sp_eeg.

## Code availability

All analysis code is available at https://github.com/sappelhoff/sp_code. The experiment presentation code is available on Zenodo (https://doi.org/10.5281/zenodo.3354368).

## Ethics information

The study was approved by the ethics committee of the Max Planck Institute for Human Development.

## Acknowledgements

We thank Agnessa Karapetian, Clara Wicharz, Jann Wäscher, Yoonsang Lee, and Zhiqi Kang for help with data collection, Dirk Ostwald and Casper Kerrén for helpful discussions and feedback, and Susannah Goss for editorial assistance.

## Author contributions

SA, RH, BS: Conceptualization, Project Administration, Writing - review & editing
SA, BS: Methodology, Writing - original draft
SA: Formal analysis, Investigation, Validation, Visualization, Data curation, Software
BS: Supervision
RH: Resources

